# Complete Genome Sequencing and Molecular Characterization of Indigenous Enterovirus A71 Genogroups D and G of India

**DOI:** 10.64898/2025.12.04.692271

**Authors:** Abhinendra Kumar, Anita Shete, Swapnil Y Varose, Santosh Jadhav, Savita Yadav, Juhi Khurana, Nivedita Gupta, Pragya D Yadav, Sarah Cherian, Madhu C Mohanty

## Abstract

Enterovirus A71 (EV-A71), a neurotropic member of the *Picornaviridae* family, comprises seven recognized genogroups (A–G). Despite prior identification of D and G genogroups in India, comprehensive full-genome characterization of these indigenous lineages remains unreported. We performed next-generation sequencing (NGS) and molecular phylogenetic analyses on two Indian isolates: R13223-IND-02 (genogroup D) and V11-2209-01 (genogroup G). Phylogenetic reconstruction revealed robust clustering of these strains within distinct clades, divergent from globally prevalent genogroups B and C, considering bootstrap support ≥99%. Both D and G genogroups appear geographically restricted to India, while genogroup D persists as an endemic lineage. Comparative genomic analyses demonstrated extensive nucleotide divergence, including numerous non-synonymous substitutions across both structural (VP1–VP4) and non-structural (2A–3D) coding regions. These findings highlight the evolutionary independence and sustained circulation of D and G genogroups in India, suggesting a unique regional evolutionary trajectory. The findings hold significant global relevance, offering valuable insights for molecular surveillance, advancing pathogenesis research, and supporting the development of genogroup-specific vaccine strategies tailored to diverse epidemiological landscapes.

## Introduction

Enterovirus A71 (EV-A71) is a causative agent of Hand, Foot, and Mouth Disease (HFMD), which has led to substantial morbidity and mortality in various Southeast Asian countries with mild infections. However, this virus has also been known to cause severe neurological manifestations including aseptic meningitis, polio-like paresis and possibly fatal encephalitis (1–3). Since the first isolation in California, USA in 1969, (4), EV-A71 associated outbreaks have been reported worldwide (5, 6). In recent years, it has gained more attention due to increasing prevalence of EV-A71 in Asia (7).

EV-A71, a non-enveloped virus, classified under the genus *Enterovirus*, within the *Picornaviridae* family, contains a ∼7.4 kb positive-sense RNA genome. This genome features a single open reading frame (ORF) with a 5′-untranslated region (UTR), a 3′-UTR and a poly(A) tail (8). The ORF translates a large polyprotein, consisting of three regions: P1 containing capsid proteins VP1-VP4; P2 and P3 containing non-structural proteins (2A-2C) and (3A - 3D) respectively (9). The VP1 protein is the outermost capsid component and serves as the major antigenic determinant (10). VP2 which is also antigenic, plays a role in virus-host interaction and EV-A71 virulence (11). The non-structural proteins, P2 and P3 are vital for replication and manipulation of host cell processes (12). The 2A protein disrupts host protein synthesis machinery, redirecting resources toward viral protein production. The primary function of 2C protein is, it works as NTPase, also participates in the replication complex assembly and interferes with innate immune defences through modulation of NF-κB signalling (13–15). The 3A protein facilitates protein–protein interactions via its N-terminal region, while 3C contributes to viral propagation by suppressing innate immunity and inducing apoptosis (16). The 3D RNA-dependent RNA polymerase (RdRp) is indispensable for genome synthesis and elongation during the replication cycle (17).

Seven genogroups (A–G) and 14 sub genogroups have been detected since the establishment of the molecular typing method of EV-A71 genogroup and sub genogroup based on entire VP1 sequences (18). Among the seven genogroups that have been delineated, B genogroup has been further divided into sub-genogroups B0–B7 and C genogroup has been divided into sub-genogroups C1–C6 (19, 20). C1 and C2 sub-genotypes are commonly found in Europe and the Asia–Pacific (21), B5 and C4 dominate in Taiwan (22), sub-genogroups D and G are confined to India (18), while E is found in Africa and F circulates in Madagascar (23).

We have identified circulating EV-A71 strains isolated from patients diagnosed with HFMD, encephalitis, and acute flaccid paralysis (AFP) (18). Two of these isolates among these, belong to indigenous genogroups D and G, which were unique to India and have not been observed in other countries. VP-1 based sequencing provided preliminary characterisation of these viruses, but their complete genomes are yet to be available publicly. To provide a comprehensive genomic profile, we performed full-length sequencing of the two indigenous EV-A71 genogroups and the circulating C genogroup. These sequences were then compared with the available sequences of EV-A71 pathogenic strains from India and neighbouring countries. Our findings seek to enhance the understanding of the genetic diversity and pathogenic potential of EV-A71 strains circulating across India.

## Methodology

The isolates of D and G genogroups grown in RD cells and characterized based on VP1 sequences were availed from ICMR-NIV, Mumbai repository and were used further for full genome sequencing and molecular characterization. Originally, both these viruses were isolated from faecal samples of AFP cases collected in the year 2001 and 2011 for D and G genogroups respectively through the WHO AFP surveillance network cases (18). NPEV serotype identification was done using partial sequencing of VP1 as previously described (18).

Similarly, the circulating C1 sub-genogroup virus was acquired from ICMR-NIV, Mumbai repository for complete genome sequencing and compared with indigenous genogroups.

The complete genome sequencing was performed by next generation sequencing (NGS) for the D genogroup isolate R13223-IND-02 and G genogroup isolate V11-2209-01. RNA extraction was done using Kingfisher flex RNA extractor (Thermofisher scientific) followed by depletion of ribosomal RNA. In the rRNA depletion step rRNA that annealed to their complimentary oligonucleotides (present in the probe hybridization buffer) were digested by thermostable RNAse H, followed by removal of genomic DNA and remaining probe with DNASe I treatment. Further, the total RNA was used for library preparation by using truesq LT library preparation kit. RNA libraries were prepared using TruSeq® Stranded mRNA Library Prep Kit. The amplified RNA libraries were quantified (Qubit™ dsDNA high sensitivity assay kit, Thermo Fisher Scientific), normalized and loaded on the Illumina sequencing MiniSeq platform. Fastp (version 0.23.4) was used to process raw paired-end reads which performed adapter trimming, quality filtering, per-read quality correction, and generated comprehensive quality control reports. *De novo* transcriptome assembly was conducted using Trinity version 2.15.2 (24) with default parameters. For phylogenetic analysis, representative full length genomes of various genogroups were downloaded from the NCBI GenBank database. Multiple sequence alignment of nucleotide sequences was carried out using the MAFFT software package (MAFFT v7.490) (25). Phylogenetic relationships were concluded using the Neighbor-Joining method. Evolutionary distances were obtained using the Maximum Composite Likelihood method and expressed as the number of base substitutions per site. Any positions containing gaps and missing data were handled using pairwise deletion option. Phylogenetic analyses were obtained using MEGA6 software (26). The amino acid mutations were analysed in the aligned file. Common and unique mutations for the genogroups of interest were obtained by comparative analysis.

## Results

The de novo assembled sequences were analysed to determine the phylogenetic lineage of the Indian EV-A71 isolates. The assembled full-length genomes were identified as belonging to genogroups D and G. Notably, prior to this study, only partial VP1 sequences of genogroups D and G had been reported from India (18); this work therefore provides the first complete genome sequences of both genogroups from the country.

A phylogenetic tree (Fig. 1) was constructed using the Neighbour-Joining method with representative EV-A71 sequences, and bootstrap resampling was applied to evaluate the reliability of the branching pattern. The analysis resolved the major EV-A71 genogroups (A– G) and their sub-genogroups (B0–B5, C1–C5) into distinct clusters, most of which were strongly supported by bootstrap values exceeding 90%, indicating a high level of confidence in the inferred evolutionary relationships.

**Figure 1.**
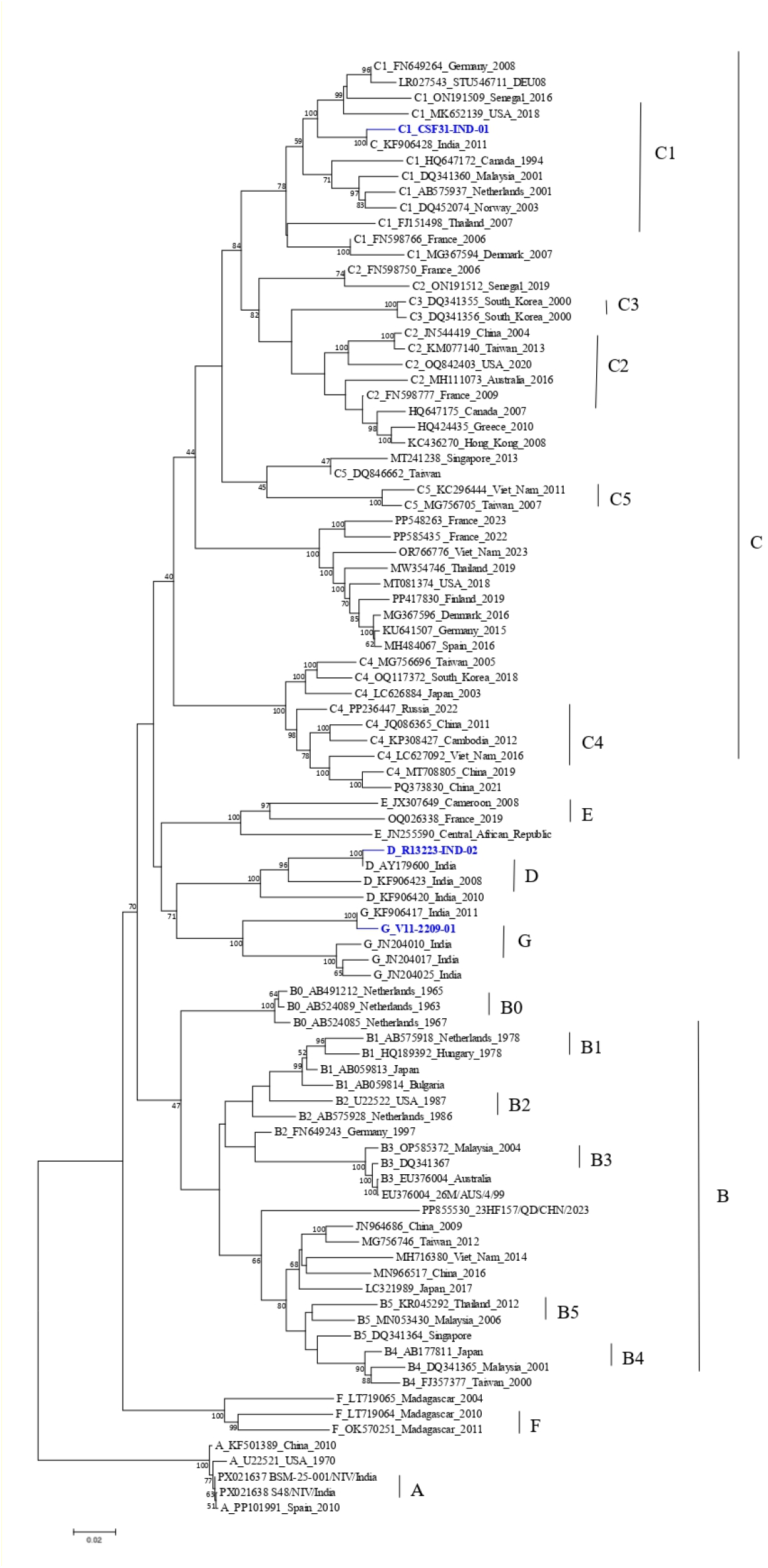
Phylogenetic tree of EV-A71 Indigenous strains using Neighbor-joining method. Neighbor-Joining (NJ) method was utilized to infer the phylogenomic relationship of indigenous strains with the representative genomes of various genogroups of EV-A71. Evolutionary distances were estimated using the Maximum Composite Likelihood method, and branch support values were estimated through 1,000 bootstrap replicates. Blue colour indicates C, D and G Genogroups of India.

The Indian strain R13223-IND-02 (NCBI accession I.D: PX260179) was placed within the genogroup D clade, clustering with earlier Indian isolates (AY179600, KF906423, KF906420) of the period 2001-2012. This grouping was supported by bootstrap value of 100%, suggesting a stable lineage and highlighting the continued circulation of genogroup D in India from the early 2000s to the present.

Likewise, the Indian strain V11-2209-01 (NCBI accession I.D: PX260180) was positioned within the genogroup G clade, forming a monophyletic group with Indian sequences of the period -2011-2012 (KF906417.1, JN204010, JN204017, and JN204025). The high bootstrap support (99%) for this cluster underscores the reliability of its placement and suggests that genogroup G represents an India-specific lineage that persists locally, in contrast to the globally widespread C and B genogroups. Overall, the well-supported clustering of genogroup D and G points to the concurrent circulation of distinct EV-A71 lineages within India, whereas the predominance of genogroup C and B in other regions highlights contrasting evolutionary trends at the global level.

A comparative evaluation of the ICMR-NIV genogroups alongside other EV-A71 genogroups showed clear patterns of both intra- and inter-genotypic divergence. Across the complete genome, the nucleotide divergence for the indigenous genogroup D with other genogroups/ sub-genogroups varied from 15.16% (B3) to 20.04% (A). Similarly, the indigenous genogroup G showed variation between 15.73% (B0) and 20.33% (A). Across the entire genome, the nucleotide divergence between C1 sub-genogroup and other genogroups/ sub-genogroups ranged from 12.57% (C3) to 19.64% (F). Across all comparisons, the amino acid identity was above 95%, which confirms the high degree of conservation of both structural and non-structural proteins despite the variability at the nucleotide level.

Comparisons of individual protein sequences, revealed notable amino acid divergence among distant genogroups up to ∼9.5% (in non-structural proteins 3B and 3C), yet VP1 remained highly conserved within sub-genogroups at levels below 2%. Among the other non-structural proteins, the 3D polymerase exhibited overall identities of more than 95%, emphasizing functional constraints on replication enzymes.

Further, the variant analysis of EV-A71 indigenous genogroups D and G was undertaken by comparing the genomes with other representative genogroup strains.

After alignment, all annotations in the multiple sequence alignment (MSA) were made with respect to the reference genotype A sequence (U22521, USA 1970). The comparison of EV-A71 genogroup D with the virulent B4 strain (DQ341365) from Malaysia (Table 1), revealed that genogroup D exhibited 66 non-synonymous and 16 synonymous mutations across the genome. The non-synonymous changes present in the various proteins were distributed as: VP1 (n=9), VP2 (n=3), VP3 (n=4), VP4 (n=2), 2A (n=4), 2B (n=2), 2C (n=8), 3A (n=2), 3B(n=2), 3C (n=8), and 3D (n=22). Synonymous mutations were observed as: VP1 (n=2), VP3 (n=1), 2A (n=3), 2B (n=1), 3A (n=2), 3C (n=3), and 3D (n=4).

**Table 1.**
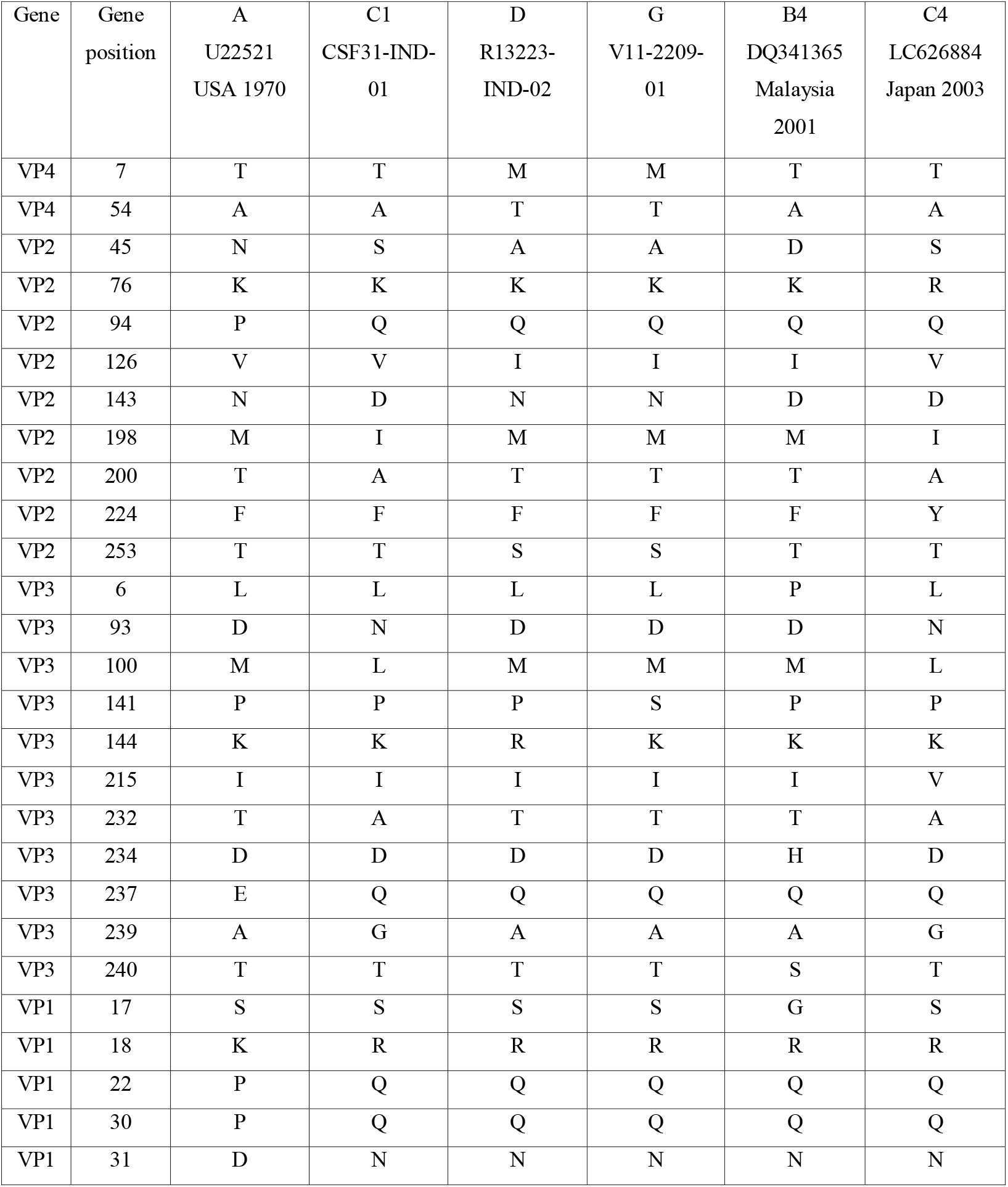
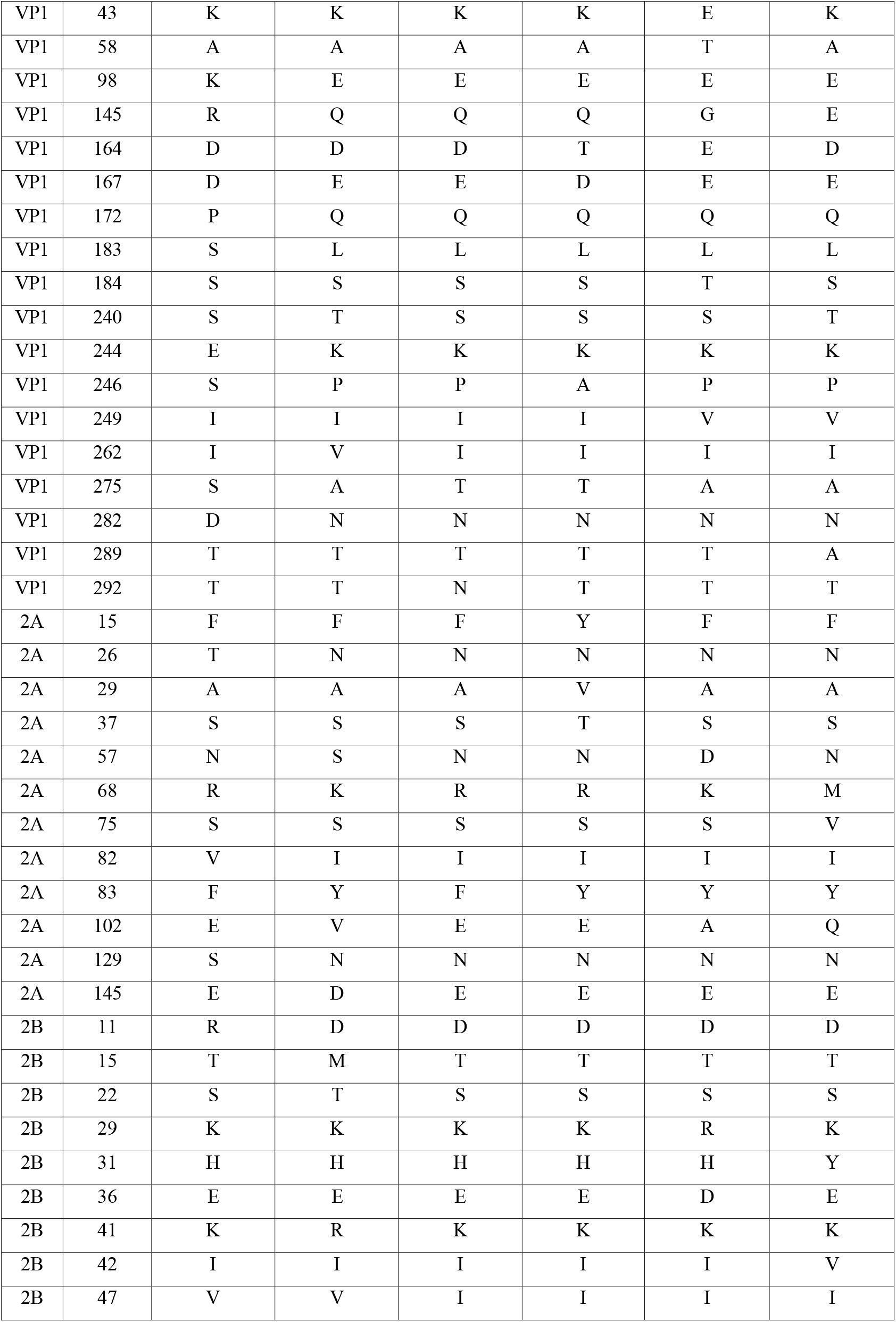
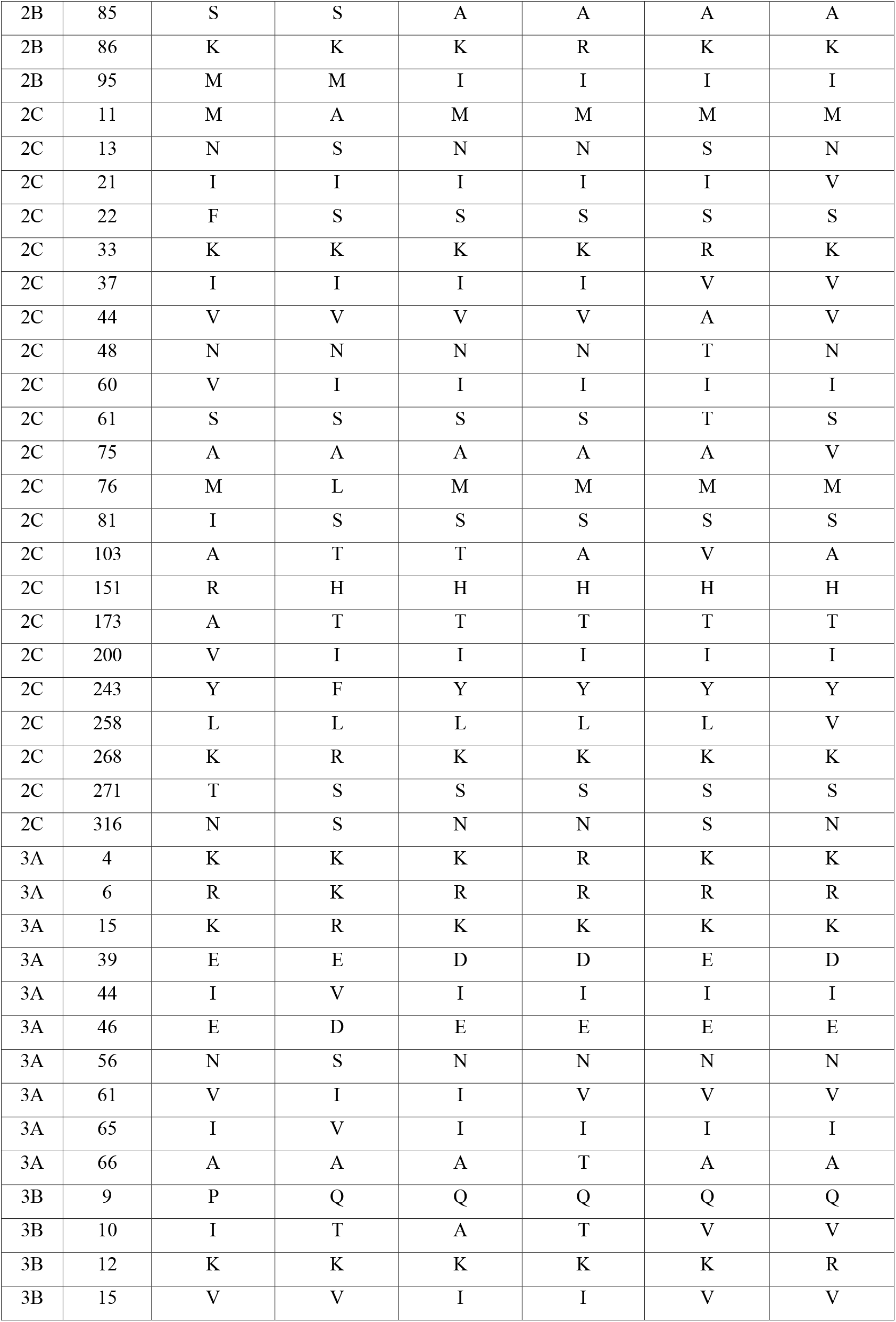
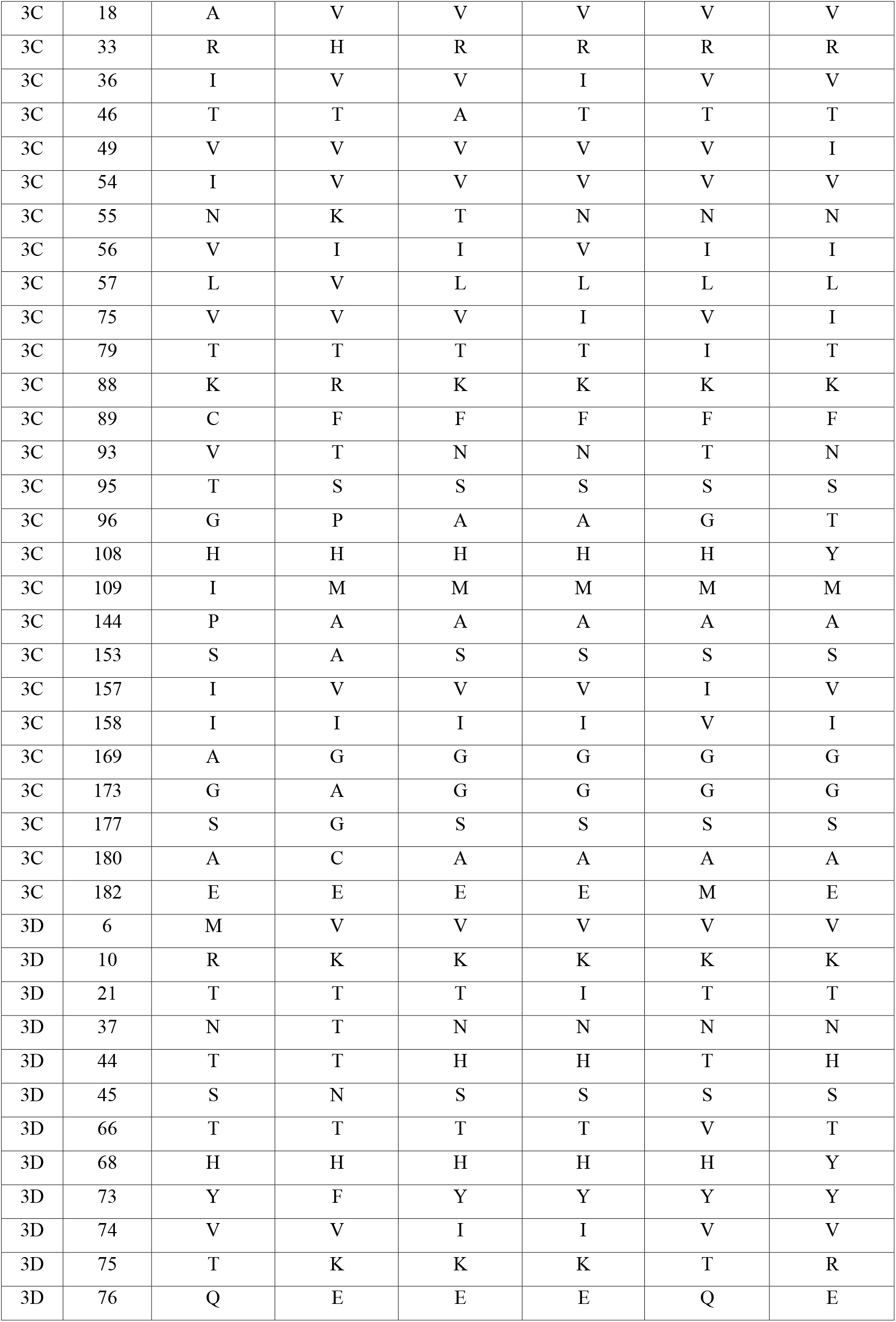
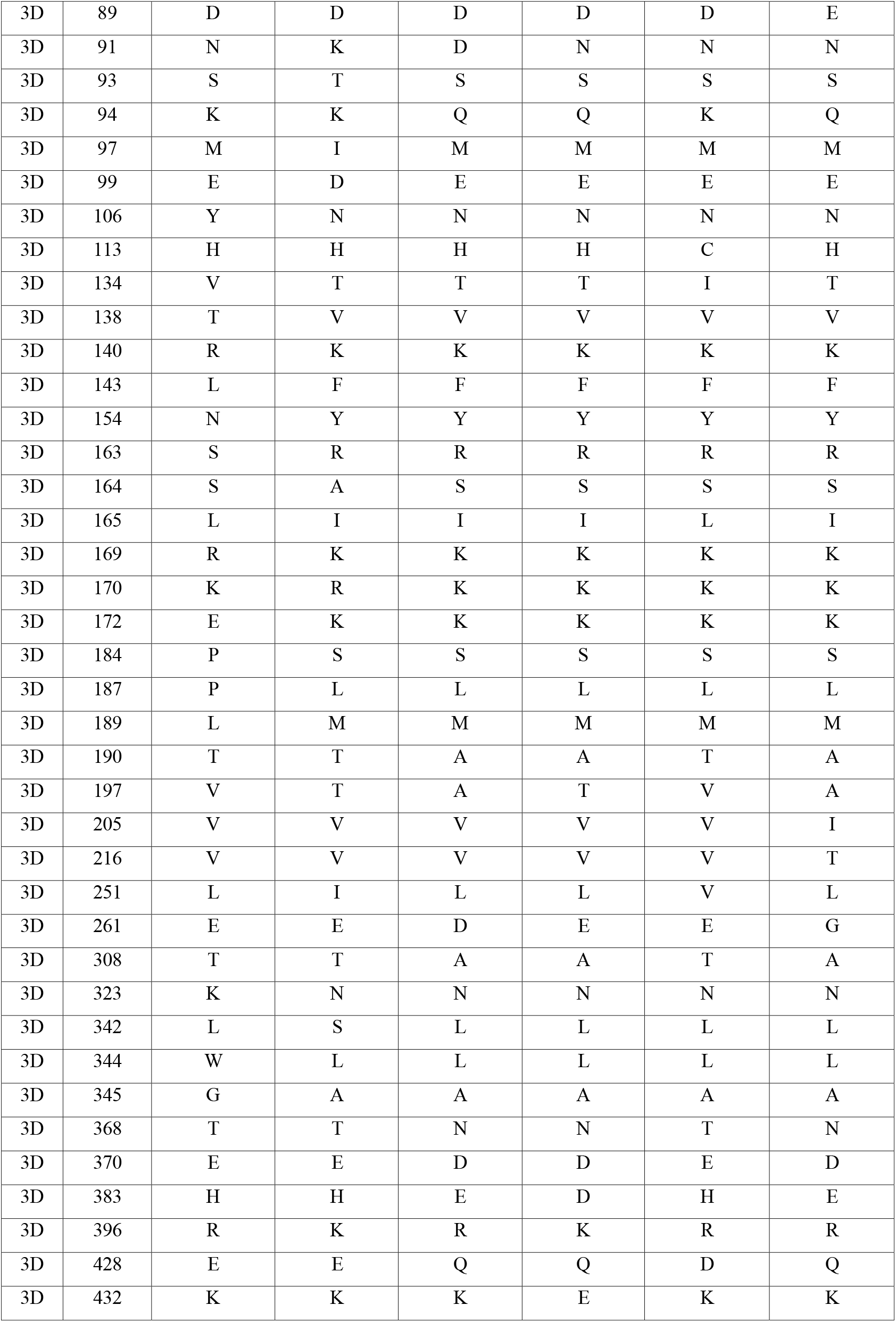
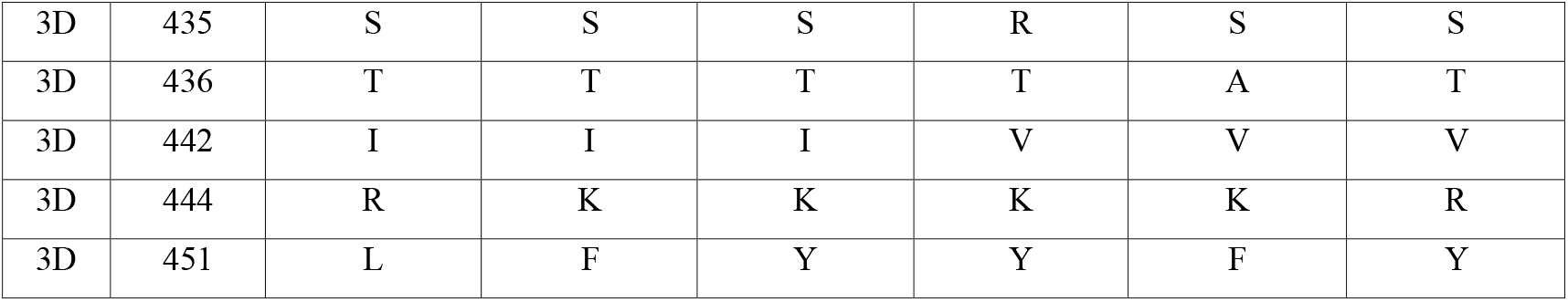
Gene-wise mutation profile of EV-A71 genogroups D and G compared with reference and neurovirulent strains. Gene-wise amino acid mutations in EV-A71 indigenous genogroups D and G were identified by comparing each gene segment (VP4, VP2, VP3, VP1, 2A, 2B, 2C, 3A, 3B, 3C, and 3D) against four representative reference strains: genogroup A (U22521, USA 1970), C1 (CSF31-IND-01), B4 (DQ341365, Malaysia, 2001), and C4 (LC626884, Japan, 2003).

The comparison of EV-A71 genogroup G with virulent B4 Malaysia Strain DQ341365, revealed a distinct mutation profile when compared with the B4 Malaysia strain, including 73 non-synonymous and 9 synonymous changes. Non-synonymous mutations were identified in VP1 (n=10), VP2 (n=3), VP3 (n=4), VP4 (n=2), 2A (n=6), 2B (n=3), 2C (n=8), 3A (n=3), 3B (n=2), 3C (n=9), and 3D (n=23). Synonymous changes were detected in VP3 (n=1), VP1 (n=1), 2A (n=2), 3C (n=2), and 3D (n=3).

Comparison of the complete genome of EV-A71 genogroup D with a virulent C4 strain (LC626884) from Japan, revealed multiple nucleotide variations compared to the strain LC626884 (Table 1). Non-synonymous amino-acid mutations identified in the structural proteins were 22, distributed as: VP1 (6), VP2 (8), VP3 (6) and VP4 (2). There were 32 non-synonymous amino-acid substitutions in the non-structural proteins as: 2A (4), 2B (3), 2C (5), 3A (1), 3B (3), 3C (6), and 3D (10). Synonymous substitutions (n=18) were found in the various proteins as follows: VP1 (n=3), VP3 (n=1), 2A (n=3), 2B (n=1), 3A (n=2), 3C (n=2), and 3D (n=6).

On the other hand, the comparison of EV-A71 indigenous genogroup G with the virulent C4 Japan strain LC626884 revealed 61 non-synonymous mutations and ten synonymous substitutions relative to the C4 strain. Non-synonymous substitutions were distributed as VP1 (8), VP2 (8), VP3 (6), VP4 (2), 2A (6), 2B (3), 2C (4), 3A (2), 3B (3), 3C (5), and 3D (14). Synonymous mutations were noted in VP1 (n=1), VP3 (n=1), 2A (n=1), 2C (n=1), 3A (n=1), 3C (n=3), and 3D (n=2).

Genogroup C has consistently been the predominant strain of EV-A71 in several Asian countries and is frequently linked to widespread outbreaks of HFMD. This genogroup is further classified into sub-genogroups C1 through C5. The comparison of EV-A71 genogroup D with sub-genogroup C1 (CSF31-IND-01) (Table-1), circulating in Asia, revealed a total of 78 non-synonymous amino acid mutations, with no synonymous substitutions detected. These amino acid changes are distributed across various viral proteins, including VP1 (n=4), VP2 (n=6), VP3 (n=5), VP4 (n=2), 2A (n=5), 2B (n=6), 2C (n=6), 3A (n=7), 3B (n=2), 3C (n=11) and 3D (n=24). On the other hand, the comparison of EV-A71 genogroup G with genogroup C1 (CSF31-IND-01) showed 89 non-synonymous mutations and 08 synonymous substitutions. The non-synonymous mutations were shown by VP1 (n=6), VP2 (n=6), VP3 (n=5), VP4 (n=2), 2A (n=7), 2B (n=7), 2C (n=7), 3A (n=10), 3B (n=1), 3C (n=13) & 3D (n=25) while synonymous mutations were observed in VP3 (n=1), VP1 (n=1), 2A (n=1), 2C (n=1), 3A (n=1), 3C (n=1), 3D (n=2).

Overall, the comparative genomic and mutational analysis of the retrieved sequences of EV-A71 genogroups D and G revealed extensive non-synonymous nucleotide variation across both structural (VP1-VP4) & non-structural (2A-3D) regions with respect to the reference strains of genogroups B4, C4 and C1. Relative to the B4 strain, both D and G genogroups exhibited extensive divergence, with 66 and 73 non-synonymous mutations, respectively. Thus, genogroup G accumulated more substitutions than D, suggesting greater evolutionary distance from the ancestral B4 lineage. Functional correlations of VP1 substitutions (G145Q, E164D/T, T184S, A275T) revealed overlap with known virulence-associated positions, while changes in 2C and 3D point to possible adaptive modifications (27). Functional correlations of the non-synonymous substitutions in the genogroup D versus C4 in the capsid protein VP1 identified changes at key antigenic sites (E145Q, T240S and V249I). Genogroup G carried a broader set of amino acid changes with respect to C4, notably in VP1 (E145Q, D164T, E167D, P246A) and multiple substitutions in 3Dpol, suggesting both antigenic drift and potential effects on polymerase function. Additionally, relative to the circulating sub-genogroup C1 (CSF31-IND-01, India), genogroup D harboured 78 non-synonymous changes. Genogroup G showed 22 non-synonymous and 8 synonymous mutations, particularly in VP1, 2A, 2C, and 3D. Several shared mutations between D and G with respect to B4, C4 and C1 (VP4 T7M, VP2 D143N, VP1 T240S, VP1 A275T, multiple 3D substitutions) (Table-1) indicate conserved evolutionary pressures across Indian lineages.

## Discussion

EV-A71, the causative agent of HFMD, is recognized as a globally significant human pathogen. Its epidemiological patterns and pathogenic mechanisms have been the subject of extensive research. Precise genotyping and classification of clinical isolates are essential for monitoring lineage circulation and emergence, as well as for understanding the determinants of severe clinical manifestations.

The HFMD was first documented in Shanghai (China) in 1981, and EV-A71 was subsequently isolated during a 2007 Linyi outbreak in Shandong Province (20). Phylogenetic and molecular epidemiology studies have confirmed the endemic presence of the C4 sub-genogroup, which is further divided to C4a and C4b (28). C4b predominated prior to 2004, after which C4a became the dominant form. Following large-scale outbreaks during 2007–2008, the Chinese Ministry of Health designated HFMD as a Category C notifiable disease on 2 May 2008. In the Asia–Pacific region, genogroups B4, B5, and C4 are most commonly observed, while genogroups C1 and C2 are more frequently found in Europe (1). Sub-genogroup C1 was first detected in Germany in 2015 and was subsequently linked to outbreaks of severe neurological illness in France, Poland, and Spain. From 2018 to early 2019, Taiwan reported HFMD cases and severe neurological complications in children infected with sub-genogroup C1 (20). Before 2000, the EV-A71 sub-genogroup C1 was mainly limited to the Western Pacific, Europe, and the United States. Subsequently, from 2000 onwards, it was becoming more dominant in Southeast Asian nations such as Malaysia, Thailand, Hong Kong, and China. B4 strains were found to be endemic in Taiwan. Molecular research shows that the VP1-A289T mutation in B4 strains diminishes CNS infectivity and neurovirulence by disrupting the VP1–vimentin binding, and VP2-K149I in C4 viruses is linked with serious neurological illness and increased virulence (29).

EV-A71 was initially isolated in India in 2001 from an AFP patient (10). Genogroups D and G have since been isolated from sporadic cases throughout India, but no documented outbreaks have been reported. In 2011–2012, sub-genogroup C1 strains were detected with AFP, HFMD and encephalitis (18). Phylogenetic analysis demonstrated that these C1 isolates were highly similar to German, Dutch, and Azerbaijani strains and implied potential epidemiological connection between Indian and European lineages (30).

We are first to report the complete genomes of Indian EV-A71 genogroups D and G. Earlier reports were limited to partial sequences (18, 29). Through NGS, we sequenced the complete genomes and aligned them with Indian C1, Malaysian B4, and Japanese C4 reference strains. Phylogenetic analysis revealed that isolates R13223-IND-02 (D) and V11-2209-01 (G) represented separate clades, distinct from internationally prevalent genogroups B and C, in concurrence with previous research that D and G are distinct Indian lineages (31).

Comparative genomic analysis documented widespread nucleotide and amino acid substitutions within both structural (VP1–VP4) and non-structural (2A–3D) domains, in line with antigenic drift and polymerase-related adaptations (27). Overlapping substitutions, such as VP4 T7M, VP2 D143N, VP1 T240S, and VP1 A275T, imply preserved evolutionary pressure (30). A number of VP1 substitutions (G145Q, E164D/T, T184S, A275T) match residues involved in virulence in Asian HFMD outbreaks (30).

VP1 is highly exposed and surface accessible and plays a central role in the assembly of virus particles, attachment, and entry. An alanine-scanning analysis using an infectious clone of EV-A71 revealed that the substitution T75A affects binding to the receptor SCARB2, viral attachment, internalization, and uncoating (32). The circulating genotypes in India showed no change at position T75 in C1, D, and G genogroups, indicating susceptibility of these genogroups to SCRAB2 receptor. Further, 145Q/G and 244K were present in all three genotypes, which have been associated with the virus-binding ability to another receptor, PSGL-1, present on leukocytes (33). According to Tan et al., 2017, residues K242 and K244 are among the factors responsible for binding to the heparan sulfate receptor observed in our D and G genogroups (34).

Evolutionary studies have demonstrated that amino acid position 145 is under positive selection and plays a major role in binding and virulence. Importantly, the 145Q substitution occurred in both the indigenous genogroups and the C1 lineage circulating in India and was reported to be a key determinant for increased infectivity in human airway organoids. However, another study (27) showed 145Q along with 164E more prevalent in severe cases, of which 164E was not observed in D and G genotype. Furthermore, 145E, 98 K/E reported to be responsible for the development of viremia and neuropathogenesis, and increased levels of cytokines in Cynomolgus monkey model, was not observed in our indigenous D and G genotypes (37). One of the major finding is 2A-68K which is linked to higher prevalence in severe cases of EV-A71 (31) and found in B4 genogroup has been observed in the circulating C1 sub-genogroup of India. Additionally, A289T which is associated with decreased viral tropism, viral genome replication, viral protein synthesis, and virus particle secretion in human brain microvascular endothelial cells (HBMECs) and decreased morbidity along with decreased CNS tropism in 1-week-old BALB/c mice has been observed to be present in both D and G genotypes indicating reduced viral attachment, infectivity in these genotypes (19).

Globally, C4 strains have taken over in China, while B4/B5 have had control in Southeast Asia and Malaysia. On the other hand, persistence of G and D in India suggests localized evolution, possibly by ecological or immunological bottlenecks. Genomic surveillance is vital to follow adaptive mutations capable of changing virulence and transmissibility, as seen in Chinese C4 strains (29).

## Conclusion

This study documents the sustained circulation of EV-A71 genogroups D and G in India. Strong phylogenetic clustering and extensive divergence from reference strains of genogroups B and C confirm their evolutionary independence and underscores regional specificity. Overall, the data reinforce that Indian EV-A71 evolution follows a unique trajectory compared to global lineages, with important implications for regional public health and vaccine strategy development. The study also highlights the importance of expanding genomic surveillance and functional characterization of Indian isolates. Assessing adaptive mutations in strains will aid vaccine development by predicting virulence and anticipating outbreak risks.

## Supporting information

E:LabDr. AnitaEV-71ASupplementary Table.docx

## Ethical Approval

The study was approved by Institutional Ethics Committee (IEC) of ICMR-NIV (Project No. NIV/IEC/Feb/2024/D-6).

## Data Availability

The authors declare that the data supporting the findings of this study is available in the article and supplementary material.

## Declaration of Generative AI and AI-assisted technologies in the writing process

No Artificial intelligence tool was used for acquiring, analyzing, interpretation of the data and drafting of the manuscript.

## Acknowledgments

The authors acknowledge Dr. Naveen Kumar (Director, ICMR-National Institute of Virology, Pune) for the constant support and guidance.

## References

1. Hagiwara A, Yoneyama T, Takami S, Hashimoto I. Genetic and phenotypic characteristics of enterovirus 71 isolates from patients with encephalitis and with hand, foot and mouth disease. Arch Virol. 1984;79(3–4):273–83.

2. Griffiths MJ, Ooi MH, Wong SC, Mohan A, Podin Y, Perera D, et al. In enterovirus 71 encephalitis with cardio-respiratory compromise, elevated interleukin 1β, interleukin 1 receptor antagonist, and granulocyte colony-stimulating factor levels are markers of poor prognosis. J Infect Dis. 2012 Sep;206(6):881–92.

3. Apostol LN, Shimizu H, Suzuki A, Umami RN, Jiao MMA, Tandoc A 3rd, et al. Molecular characterization of enterovirus-A71 in children with acute flaccid paralysis in the Philippines. BMC Infect Dis. 2019 May;19(1):370.

4. Blomberg J, Lycke E, Ahlfors K, Johnsson T, Wolontis S, von Zeipel G. Letter: New enterovirus type associated with epidemic of aseptic meningitis and-or hand, foot, and mouth disease. Lancet (London, England). 1974 Jul;2(7872):112.

5. Schuffenecker I, Mirand A, Antona D, Henquell C, Chomel JJ, Archimbaud C, et al. Epidemiology of human enterovirus 71 infections in France, 2000-2009. J Clin Virol Off Publ Pan Am Soc Clin Virol. 2011 Jan;50(1):50–6.

6. McMinn P, Stratov I, Nagarajan L, Davis S. Neurological manifestations of enterovirus 71 infection in children during an outbreak of hand, foot, and mouth disease in Western Australia. Clin Infect Dis an Off Publ Infect Dis Soc Am. 2001 Jan;32(2):236–42.

7. Komatsu H, Shimizu Y, Takeuchi Y, Ishiko H, Takada H. Outbreak of severe neurologic involvement associated with Enterovirus 71 infection. Pediatr Neurol. 1999 Jan;20(1):17–23.

8. Yuan J, Shen L, Wu J, Zou X, Gu J, Chen J, et al. Enterovirus A71 Proteins: Structure and Function. Front Microbiol. 2018;9:286.

9. Shen M, Reitman ZJ, Zhao Y, Moustafa I, Wang Q, Arnold JJ, et al. Picornavirus Genome Replication: IDENTIFICATION OF THE SURFACE OF THE POLIOVIRUS (PV) 3C DIMER THAT INTERACTS WITH PV 3Dpol DURING VPg URIDYLYLATION AND CONSTRUCTION OF A STRUCTURAL MODEL FOR THE PV 3C2-3Dpol COMPLEX*. J Biol Chem [Internet]. 2008;283(2):875–88. Available from: https://www.sciencedirect.com/science/article/pii/S0021925820689980

10. Keng TK, Tsan-Yuk LT, Fun CY, M. BJ, Adeeba K, William TCY, et al. Evolutionary Genetics of Human Enterovirus 71: Origin, Population Dynamics, Natural Selection, and Seasonal Periodicity of the VP1 Gene. J Virol [Internet]. 2010 Apr 1;84(7):3339–50. Available from: 10.1128/jvi.01019-09

11. Kiener TK, Jia Q, Lim XF, He F, Meng T, Kwong Chow VT, et al. Characterization and specificity of the linear epitope of the enterovirus 71 VP2 protein. Virol J [Internet]. 2012;9(1):55. Available from: 10.1186/1743-422X-9-55

12. Huang PN, Shih SR. Update on enterovirus 71 infection. Curr Opin Virol. 2014 Apr;5:98–104.

13. Li Z, Ning S, Su X, Liu X, Wang H, Liu Y, et al. Enterovirus 71 antagonizes the inhibition of the host intrinsic antiviral factor A3G. Nucleic Acids Res. 2018 Nov;46(21):11514–27.

14. Zheng Z, Li H, Zhang Z, Meng J, Mao D, Bai B, et al. Enterovirus 71 2C Protein Inhibits TNF-α–Mediated Activation of NF-κB by Suppressing IκB Kinase β Phosphorylation. J Immunol. 2011 Sep;187(5):2202–12.

15. Du H, Yin P, Yang X, Zhang L, Jin Q, Zhu G. Enterovirus 71 2C Protein Inhibits NF-κB Activation by Binding to RelA(p65). Sci Rep. 2015 Sep;5:14302.

16. Bai Y, Yao L, Wei T, Tian F, Jin DY, Chen L, et al. Presumed Asymptomatic Carrier Transmission of COVID-19. JAMA. 2020 Apr;323(14):1406–7.

17. Diarimalala RO, Hu M, Wei Y, Hu K. Recent advances of enterovirus 71 3Cpro targeting Inhibitors. Virol J. 2020 Nov;17(1):173.

18. Saxena VK, Sane S, Nadkarni SS, Sharma DK, Deshpande JM. Genetic diversity of enterovirus A71, India. Emerg Infect Dis. 2015 Jan;21(1):123–6.

19. Ang PY, Chong CWH, Alonso S. Viral determinants that drive Enterovirus-A71 fitness and virulence. Emerg Microbes Infect. 2021 Dec;10(1):713–24.

20. Oberste MS, Maher K, Kilpatrick DR, Flemister MR, Brown BA, Pallansch MA. Typing of human enteroviruses by partial sequencing of VP1. J Clin Microbiol. 1999 May;37(5):1288–93.

21. Van Tu P, Thao NTT, Perera D, Truong KH, Tien NTK, Thuong TC, et al. Epidemiologic and virologic investigation of hand, foot, and mouth disease, southern Vietnam, 2005. Emerg Infect Dis. 2007 Nov;13(11):1733–41.

22. Huang YP, Lin TL, Kuo CY, Lin MW, Yao CY, Liao HW, et al. The circulation of subgenogroups B5 and C5 of enterovirus 71 in Taiwan from 2006 to 2007. Virus Res. 2008;137(2):206–12.

23. Fernandez-Garcia MD, Volle R, Joffret ML, Sadeuh-Mba SA, Gouandjika-Vasilache I, Kebe O, et al. Genetic Characterization of Enterovirus A71 Circulating in Africa. Emerg Infect Dis. 2018 Apr;24(4):754–7.

24. Grabherr MG, Haas BJ, Yassour M, Levin JZ, Thompson DA, Amit I, et al. Fulllength transcriptome assembly from RNA-Seq data without a reference genome. Nat Biotechnol. 2011 May;29(7):644–52.

25. Katoh K, Standley DM. MAFFT multiple sequence alignment software version 7: improvements in performance and usability. Mol Biol Evol. 2013 Apr;30(4):772–80.

26. Tamura K, Stecher G, Peterson D, Filipski A, Kumar S. MEGA6: Molecular Evolutionary Genetics Analysis version 6.0. Mol Biol Evol. 2013 Dec;30(12):2725–9.

27. Sheng-Wen H, Ching-Hui T, M. FJ, Chin-Hui L, Shih-Min W, Ching-Chung L, et al. Mapping Enterovirus A71 Antigenic Determinants from Viral Evolution. J Virol [Internet]. 2015 Oct 22;89(22):11500–6. Available from: 10.1128/jvi.02035-15

28. Thao NTT, Donato C, Trang VTH, Kien NT, Trang PMT, Khanh TQ, et al. Evolution and Spatiotemporal Dynamics of Enterovirus A71 Subgenogroups in Vietnam. J Infect Dis [Internet]. 2017 Dec 12;216(11):1371–9. Available from: 10.1093/infdis/jix500

29. Puenpa J, Wanlapakorn N, Vongpunsawad S, Poovorawan Y. The History of Enterovirus A71 Outbreaks and Molecular Epidemiology in the Asia-Pacific Region. J Biomed Sci [Internet]. 2019;26(1):75. Available from: 10.1186/s12929-019-0573-2

30. Zhu Y, Songya L, Jianchun X, Jun Y, Li X, Jun L, et al. Complete Genome Sequence of a Human Enterovirus 71 Strain Isolated in Wuhan, China, in 2010. Genome Announc [Internet]. 2013 Dec 26;1(6):10.1128/genomea.01112-13. Available from: 10.1128/genomea.01112-13

31. Li R, Zou Q, Chen L, Zhang H, Wang Y. Molecular analysis of virulent determinants of enterovirus 71. PLoS One. 2011;6(10):e26237.

32. Zhang W, Li Q, Yi D, Zheng R, Liu G, Liu Q, et al. Novel virulence determinants in VP1 regulate the assembly of enterovirus-A71. J Virol. 2024 Dec;98(12):e0165524.

33. Nishimura Y, Lee H, Hafenstein S, Kataoka C, Wakita T, Bergelson JM, et al. Enterovirus 71 Binding to PSGL-1 on Leukocytes: VP1-145 Acts as a Molecular Switch to Control Receptor Interaction. PLOS Pathog [Internet]. 2013 Jul 25;9(7):e1003511. Available from: 10.1371/journal.ppat.1003511

34. Tan CW, Sam IC, Lee VS, Wong HV, Chan YF. VP1 residues around the five-fold axis of enterovirus A71 mediate heparan sulfate interaction. Virology. 2017 Jan;501:79–87.

35. Keng TK, Tsan-Yuk LT, Fun CY, M. BJ, Adeeba K, William TCY, et al. Evolutionary Genetics of Human Enterovirus 71: Origin, Population Dynamics, Natural Selection, and Seasonal Periodicity of the VP1 Gene. J Virol. 2010 Apr;84(7):3339–50.

36. van der Sanden SMG, Sachs N, Koekkoek SM, Koen G, Pajkrt D, Clevers H, et al. Enterovirus 71 infection of human airway organoids reveals VP1-145 as a viral infectivity determinant. Emerg Microbes Infect. 2018 May;7(1):84.

37. Kataoka C, Suzuki T, Kotani O, Iwata-Yoshikawa N, Nagata N, Ami Y, et al. The Role of VP1 Amino Acid Residue 145 of Enterovirus 71 in Viral Fitness and Pathogenesis in a Cynomolgus Monkey Model. PLOS Pathog [Internet]. 2015 Jul 16;11(7):e1005033. Available from: 10.1371/journal.ppat.1005033

