## Supplementary material for "Complete Genome Sequencing and Molecular Characterization of Indigenous Enterovirus A71 Genogroups D and G of India": E:LabDr. AnitaEV-71ASupplementary Table.docx

Supplementary Table-1: Nucleotide: Gene wise Comparison of p-distance (NIV vs other)

|  | Genogroup | 5'UTR | VP4(1A) | VP2 | VP3 | VP1 | 2A | 2B | 2C | 3A | 3B | 3C | 3D | 3'UTR | Overall |
| --- | --- | --- | --- | --- | --- | --- | --- | --- | --- | --- | --- | --- | --- | --- | --- |
| NIV-C vs | A | 0.1519 | 0.2006 | 0.1834 | 0.174 | 0.1633 | 0.2035 | 0.217 | 0.1983 | 0.232 | 0.2677 | 0.2307 | 0.2124 | 0.1165 | 0.1932 |
|  | B0 |  |  |  |  | 0.1641 |  |  |  |  |  |  |  |  | 0.1641 |
|  | B1 | 0.1304 | 0.1693 | 0.1621 | 0.177 | 0.1703 | 0.2002 | 0.2645 | 0.1999 | 0.2557 | 0.2194 | 0.234 | 0.1958 | 0.1304 | 0.1794 |
|  | B2 | 0.1134 | 0.1475 | 0.1558 | 0.1722 | 0.1634 | 0.1858 | 0.2477 | 0.2073 | 0.2654 | 0.2045 | 0.2372 | 0.2081 | 0.1255 | 0.1787 |
|  | B3 | 0.1074 | 0.142 | 0.1726 | 0.1645 | 0.1619 | 0.1828 | 0.2276 | 0.2028 | 0.2546 | 0.1831 | 0.2429 | 0.2073 | 0.1614 | 0.1846 |
|  | B4 | 0.1228 | 0.1551 | 0.1831 | 0.1702 | 0.1774 | 0.2047 | 0.2619 | 0.2015 | 0.2441 | 0.2121 | 0.2476 | 0.2027 | 0.0783 | 0.1872 |
|  | B5 | 0.1214 | 0.1644 | 0.181 | 0.1761 | 0.183 | 0.1894 | 0.2547 | 0.2095 | 0.2524 | 0.1951 | 0.2512 | 0.2058 | 0.1 | 0.1935 |
|  | C1 | 0.1248 | 0.0853 | 0.0722 | 0.0678 | 0.0623 | 0.0793 | 0.0895 | 0.0799 | 0.102 | 0.0756 | 0.0763 | 0.068 | 0.0396 | 0.0708 |
|  | C2 | 0.1586 | 0.1391 | 0.125 | 0.1287 | 0.1083 | 0.1569 | 0.1532 | 0.1387 | 0.152 | 0.111 | 0.1278 | 0.1272 | 0.0693 | 0.1284 |
|  | C3 | 0.1591 | 0.1014 | 0.1257 | 0.1226 | 0.1055 | 0.1578 | 0.1316 | 0.1142 | 0.1282 | 0.0836 | 0.112 | 0.1353 | 0.0361 | 0.1257 |
|  | C4 | 0.13 | 0.1321 | 0.1245 | 0.1258 | 0.1239 | 0.1536 | 0.249 | 0.1953 | 0.2642 | 0.2262 | 0.2473 | 0.2225 | 0.1781 | 0.1754 |
|  | C5 | 0.1604 | 0.0845 | 0.134 | 0.106 | 0.1234 | 0.1368 | 0.1462 | 0.1524 | 0.198 | 0.1512 | 0.152 | 0.1696 | 0.0836 | 0.1363 |
|  | D |  |  |  |  | 0.1628 |  |  |  |  |  |  |  |  | 0.1628 |
|  | E | 0.1295 | 0.1741 | 0.1707 | 0.1927 | 0.1715 | 0.1896 | 0.197 | 0.1987 | 0.2037 | 0.197 | 0.2404 | 0.192 | 0.037 | 0.1788 |
|  | F | 0.1353 | 0.158 | 0.1996 | 0.1928 | 0.1832 | 0.2132 | 0.2321 | 0.2158 | 0.2309 | 0.2374 | 0.2407 | 0.1968 | 0.0843 | 0.1964 |
|  | G |  |  |  |  | 0.1521 |  |  |  |  |  |  |  |  | 0.1521 |
|  | Un | 0.1254 | 0.1011 | 0.1001 | 0.1075 | 0.0961 | 0.1259 | 0.2087 | 0.2213 | 0.2267 | 0.2357 | 0.2399 | 0.2128 | 0.3661 | 0.1636 |
| NIV-D vs | A | 0.1636 | 0.1739 | 0.1978 | 0.181 | 0.1809 | 0.2179 | 0.1901 | 0.2111 | 0.2351 | 0.298 | 0.2368 | 0.2091 | 0.2105 | 0.2004 |
|  | B0 |  |  |  |  | 0.1757 |  |  |  |  |  |  |  |  | 0.1757 |
|  | B1 | 0.1106 | 0.1763 | 0.1594 | 0.1792 | 0.1774 | 0.217 | 0.1985 | 0.1525 | 0.1783 | 0.1515 | 0.2058 | 0.2207 | 0.2488 | 0.1795 |
|  | B2 | 0.1035 | 0.1787 | 0.1589 | 0.184 | 0.1734 | 0.218 | 0.2002 | 0.1616 | 0.1725 | 0.1288 | 0.2031 | 0.2204 | 0.2482 | 0.1769 |
|  | B3 | 0.097 | 0.1932 | 0.1816 | 0.1778 | 0.1759 | 0.2019 | 0.1726 | 0.1499 | 0.1519 | 0.1879 | 0.1224 | 0.1218 | 0.0914 | 0.1516 |
|  | B4 | 0.1124 | 0.1937 | 0.1733 | 0.1833 | 0.1753 | 0.2148 | 0.2002 | 0.1626 | 0.1628 | 0.1136 | 0.2149 | 0.2078 | 0.1914 | 0.1783 |
|  | B5 | 0.1177 | 0.1896 | 0.183 | 0.1712 | 0.174 | 0.2187 | 0.1872 | 0.1686 | 0.1667 | 0.1402 | 0.2132 | 0.2168 | 0.2299 | 0.183 |
|  | C1 | 0.1611 | 0.1836 | 0.193 | 0.1886 | 0.1655 | 0.1843 | 0.2469 | 0.1972 | 0.2297 | 0.2449 | 0.2365 | 0.2117 | 0.178 | 0.1863 |
|  | C2 | 0.1603 | 0.1643 | 0.1902 | 0.1742 | 0.1713 | 0.1944 | 0.2427 | 0.2083 | 0.2282 | 0.2083 | 0.2372 | 0.2208 | 0.1889 | 0.1922 |
|  | C3 | 0.1685 | 0.1739 | 0.2042 | 0.1571 | 0.1819 | 0.1924 | 0.2541 | 0.2042 | 0.2306 | 0.197 | 0.2423 | 0.2038 | 0.1605 | 0.1969 |
|  | C4 | 0.1342 | 0.1761 | 0.1904 | 0.1778 | 0.1718 | 0.1879 | 0.2103 | 0.1683 | 0.1714 | 0.1734 | 0.1666 | 0.1548 | 0.1091 | 0.1688 |
|  | C5 | 0.1753 | 0.1908 | 0.1871 | 0.1709 | 0.1833 | 0.2002 | 0.244 | 0.2006 | 0.2442 | 0.2348 | 0.245 | 0.2254 | 0.2873 | 0.196 |
|  | D |  |  |  |  | 0.0573 |  |  |  |  |  |  |  |  | 0.0573 |
|  | E | 0.0999 | 0.1691 | 0.1865 | 0.1823 | 0.1709 | 0.2064 | 0.187 | 0.1966 | 0.186 | 0.2879 | 0.2204 | 0.2241 | 0.1519 | 0.1826 |
|  | F | 0.1187 | 0.1787 | 0.1558 | 0.1764 | 0.1904 | 0.2261 | 0.2038 | 0.2003 | 0.1938 | 0.2727 | 0.2265 | 0.2132 | 0.1811 | 0.1898 |
|  | G |  |  |  |  | 0.1623 |  |  |  |  |  |  |  |  | 0.1623 |
|  | Un | 0.1241 | 0.1798 | 0.1935 | 0.1786 | 0.1617 | 0.171 | 0.2028 | 0.2004 | 0.1895 | 0.2576 | 0.2267 | 0.1636 | 0.1462 | 0.1781 |
| NIV-G vs | A | 0.1731 | 0.1839 | 0.1899 | 0.1706 | 0.1987 | 0.2243 | 0.2256 | 0.2073 | 0.2158 | 0.2828 | 0.2319 | 0.2188 | 0.1804 | 0.2033 |
|  | B0 |  |  |  |  | 0.1573 |  |  |  |  |  |  |  |  | 0.1573 |
|  | B1 | 0.1211 | 0.1815 | 0.166 | 0.1742 | 0.1702 | 0.2076 | 0.1852 | 0.1759 | 0.1647 | 0.1818 | 0.2049 | 0.2077 | 0.227 | 0.1754 |
|  | B2 | 0.1093 | 0.1791 | 0.1741 | 0.1742 | 0.1713 | 0.2032 | 0.1818 | 0.1694 | 0.1609 | 0.197 | 0.214 | 0.2197 | 0.2397 | 0.1768 |
|  | B3 | 0.1005 | 0.1741 | 0.1781 | 0.172 | 0.1685 | 0.1821 | 0.1481 | 0.1712 | 0.1775 | 0.2152 | 0.1497 | 0.1521 | 0.116 | 0.1593 |
|  | B4 | 0.1252 | 0.1844 | 0.1675 | 0.1687 | 0.178 | 0.2054 | 0.2054 | 0.1583 | 0.1686 | 0.2045 | 0.2113 | 0.2136 | 0.216 | 0.1792 |
|  | B5 | 0.1277 | 0.1935 | 0.1751 | 0.1696 | 0.1709 | 0.2068 | 0.1965 | 0.1669 | 0.1628 | 0.2424 | 0.2129 | 0.2178 | 0.2334 | 0.1831 |
|  | C1 | 0.1653 | 0.1518 | 0.1885 | 0.1903 | 0.1622 | 0.211 | 0.2315 | 0.1997 | 0.2456 | 0.2184 | 0.2472 | 0.203 | 0.2013 | 0.1857 |
|  | C2 | 0.1578 | 0.1927 | 0.1875 | 0.1725 | 0.1742 | 0.2151 | 0.2365 | 0.1886 | 0.2258 | 0.2216 | 0.2498 | 0.2102 | 0.2136 | 0.1915 |
|  | C3 | 0.167 | 0.1549 | 0.1754 | 0.1756 | 0.1834 | 0.2121 | 0.229 | 0.194 | 0.2248 | 0.2121 | 0.2514 | 0.2163 | 0.1852 | 0.1973 |
|  | C4 | 0.1424 | 0.1495 | 0.1899 | 0.1776 | 0.1721 | 0.2001 | 0.2144 | 0.1545 | 0.1714 | 0.2441 | 0.1858 | 0.1658 | 0.1043 | 0.1719 |
|  | C5 | 0.1773 | 0.1518 | 0.2003 | 0.1866 | 0.1653 | 0.2165 | 0.2374 | 0.1921 | 0.2054 | 0.2197 | 0.2477 | 0.2149 | 0.3532 | 0.1897 |
|  | D |  |  |  |  | 0.1493 |  |  |  |  |  |  |  |  | 0.1493 |
|  | E | 0.1122 | 0.1839 | 0.1669 | 0.1579 | 0.169 | 0.2233 | 0.2155 | 0.2011 | 0.1977 | 0.2576 | 0.2149 | 0.2158 | 0.1646 | 0.179 |
|  | F | 0.125 | 0.1916 | 0.161 | 0.1862 | 0.175 | 0.1939 | 0.2121 | 0.1988 | 0.1977 | 0.2677 | 0.2186 | 0.2193 | 0.1975 | 0.1894 |
|  | G |  |  |  |  | 0.1056 |  |  |  |  |  |  |  |  | 0.1056 |
|  | Un | 0.1215 | 0.17 | 0.1841 | 0.1841 | 0.1577 | 0.1957 | 0.217 | 0.1949 | 0.2007 | 0.2525 | 0.2242 | 0.1555 | 0.2008 | 0.1768 |

Supplementary Table-2: Amino acid: Gene wise Comparison of p-distance (NIV vs other)

| AA | Genogroup | 5'UTR | VP4(1A) | VP2 | VP3 | VP1 | 2A | 2B | 2C | 3A | 3B | 3C | 3D | 3'UTR | Overall |
| --- | --- | --- | --- | --- | --- | --- | --- | --- | --- | --- | --- | --- | --- | --- | --- |
| NIV-C vs | A |  | 0 | 0.0186 | 0.0179 | 0.0366 | 0.058 | 0.0337 | 0.0332 | 0.0814 | 0.0606 | 0.0947 | 0.0628 |  | 0.0447 |
|  | B0 |  |  |  |  | 0.0188 |  |  |  |  |  |  |  |  | 0.0188 |
|  | B1 |  | 0 | 0.0198 | 0.0248 | 0.0337 | 0.0136 | 0.0909 | 0.0337 | 0.0872 | 0.0455 | 0.0601 | 0.0487 |  | 0.0368 |
|  | B2 |  | 0 | 0.018 | 0.0289 | 0.0335 | 0.0335 | 0.0808 | 0.0398 | 0.0814 | 0.0682 | 0.0601 | 0.0509 |  | 0.039 |
|  | B3 |  | 0.0029 | 0.015 | 0.0264 | 0.0291 | 0.0242 | 0.0667 | 0.0304 | 0.0977 | 0.0455 | 0.0601 | 0.0603 |  | 0.0399 |
|  | B4 |  | 0.0074 | 0.016 | 0.031 | 0.03 | 0.0202 | 0.0808 | 0.0292 | 0.0814 | 0.0455 | 0.071 | 0.0455 |  | 0.0356 |
|  | B5 |  | 0.0018 | 0.02 | 0.0289 | 0.0282 | 0.0252 | 0.0795 | 0.0345 | 0.0799 | 0.0511 | 0.0848 | 0.053 |  | 0.0418 |
|  | C1 |  | 0.0018 | 0.0013 | 0.0015 | 0.0062 | 0.0202 | 0.0152 | 0.0154 | 0.0296 | 0.0057 | 0.0144 | 0.0175 |  | 0.0098 |
|  | C2 |  | 0.0036 | 0.0061 | 0.0052 | 0.0164 | 0.0269 | 0.0391 | 0.0283 | 0.0625 | 0.0057 | 0.0143 | 0.0315 |  | 0.0209 |
|  | C3 |  | 0 | 0.0003 | 0.0041 | 0.0072 | 0.0369 | 0.0303 | 0.0134 | 0.0581 | 0 | 0.0082 | 0.0249 |  | 0.0156 |
|  | C4 |  | 0 | 0.0095 | 0.0028 | 0.0202 | 0.0329 | 0.0752 | 0.0325 | 0.1047 | 0.0859 | 0.0747 | 0.0676 |  | 0.04 |
|  | C5 |  | 0 | 0.0023 | 0.0041 | 0.0094 | 0.0369 | 0.0455 | 0.024 | 0.0698 | 0 | 0.0301 | 0.026 |  | 0.0184 |
|  | D |  |  |  |  | 0.0153 |  |  |  |  |  |  |  |  | 0.0153 |
|  | E |  | 0 | 0.0239 | 0.0363 | 0.0323 | 0.0611 | 0.0404 | 0.017 | 0.093 | 0.0455 | 0.082 | 0.0402 |  | 0.0389 |
|  | F |  | 0.0145 | 0.0238 | 0.0193 | 0.0309 | 0.0513 | 0.0438 | 0.0259 | 0.1163 | 0.0152 | 0.0729 | 0.0541 |  | 0.0411 |
|  | G |  |  |  |  | 0.0454 |  |  |  |  |  |  |  |  | 0.0454 |
|  | Un |  | 0 | 0.0038 | 0.0092 | 0.0068 | 0.0225 | 0.0338 | 0.0248 | 0.0982 | 0.0354 | 0.0686 | 0.0548 |  | 0.0306 |
| NIV-D vs | A |  | 0.029 | 0.0145 | 0.0072 | 0.0364 | 0.0255 | 0.0337 | 0.0162 | 0.0233 | 0.1061 | 0.0619 | 0.0622 |  | 0.0343 |
|  | B0 |  |  |  |  | 0.0151 |  |  |  |  |  |  |  |  | 0.0151 |
|  | B1 |  | 0.029 | 0.0157 | 0.0127 | 0.0312 | 0.021 | 0.0303 | 0.0228 | 0.0407 | 0.0909 | 0.0301 | 0.0597 |  | 0.0321 |
|  | B2 |  | 0.029 | 0.0141 | 0.0168 | 0.0309 | 0.0377 | 0.0202 | 0.035 | 0.0233 | 0.1136 | 0.0328 | 0.0575 |  | 0.0335 |
|  | B3 |  | 0.0319 | 0.018 | 0.0202 | 0.0348 | 0.0317 | 0.0061 | 0.0134 | 0.0209 | 0.0909 | 0.023 | 0.0248 |  | 0.0233 |
|  | B4 |  | 0.0366 | 0.0121 | 0.0189 | 0.0343 | 0.0277 | 0.0202 | 0.0243 | 0.0233 | 0.0909 | 0.0437 | 0.0456 |  | 0.0323 |
|  | B5 |  | 0.0308 | 0.0161 | 0.0174 | 0.0325 | 0.0344 | 0.0215 | 0.0293 | 0.0276 | 0.0852 | 0.0588 | 0.0554 |  | 0.0354 |
|  | C1 |  | 0.029 | 0.0239 | 0.0215 | 0.0219 | 0.0335 | 0.0568 | 0.0289 | 0.0613 | 0.0852 | 0.0643 | 0.0539 |  | 0.033 |
|  | C2 |  | 0.029 | 0.0298 | 0.022 | 0.0276 | 0.026 | 0.0543 | 0.0304 | 0.048 | 0.0966 | 0.0594 | 0.0497 |  | 0.0358 |
|  | C3 |  | 0.029 | 0.0239 | 0.021 | 0.0253 | 0.0377 | 0.0505 | 0.0182 | 0.0465 | 0.0909 | 0.0683 | 0.05 |  | 0.036 |
|  | C4 |  | 0.029 | 0.0296 | 0.0228 | 0.0313 | 0.0292 | 0.0146 | 0.0155 | 0.0233 | 0.096 | 0.0376 | 0.0242 |  | 0.0262 |
|  | C5 |  | 0.029 | 0.0259 | 0.021 | 0.0253 | 0.0444 | 0.0657 | 0.0289 | 0.0465 | 0.0909 | 0.0738 | 0.0532 |  | 0.0352 |
|  | D |  |  |  |  | 0.0107 |  |  |  |  |  |  |  |  | 0.0107 |
|  | E |  | 0.029 | 0.0082 | 0.003 | 0.0298 | 0.041 | 0.0202 | 0.0061 | 0.0116 | 0.1364 | 0.0546 | 0.0479 |  | 0.0279 |
|  | F |  | 0.0338 | 0.0093 | 0.0072 | 0.0287 | 0.0255 | 0.0303 | 0.0111 | 0.0426 | 0.0909 | 0.0383 | 0.0608 |  | 0.0302 |
|  | G |  |  |  |  | 0.0425 |  |  |  |  |  |  |  |  | 0.0425 |
|  | Un |  | 0.029 | 0.0274 | 0.0178 | 0.0249 | 0.0166 | 0.0427 | 0.0145 | 0.0194 | 0.1212 | 0.0498 | 0.0242 |  | 0.0259 |
| NIV-G vs | A |  | 0.029 | 0.0146 | 0.0069 | 0.0288 | 0.0573 | 0.0438 | 0.0132 | 0.0349 | 0.1061 | 0.0455 | 0.0685 |  | 0.036 |
|  | B0 |  |  |  |  | 0.0241 |  |  |  |  |  |  |  |  | 0.0241 |
|  | B1 |  | 0.029 | 0.0157 | 0.0124 | 0.0334 | 0.0395 | 0.0404 | 0.0228 | 0.0523 | 0.0909 | 0.0355 | 0.0552 |  | 0.0344 |
|  | B2 |  | 0.029 | 0.0143 | 0.0165 | 0.0333 | 0.0562 | 0.0303 | 0.035 | 0.0349 | 0.1136 | 0.0383 | 0.0552 |  | 0.0356 |
|  | B3 |  | 0.0319 | 0.0179 | 0.0198 | 0.0332 | 0.0502 | 0.0162 | 0.0103 | 0.0279 | 0.0909 | 0.0175 | 0.0268 |  | 0.0246 |
|  | B4 |  | 0.0366 | 0.0124 | 0.0186 | 0.0299 | 0.0462 | 0.0303 | 0.0243 | 0.0349 | 0.0909 | 0.0492 | 0.0476 |  | 0.0322 |
|  | B5 |  | 0.0308 | 0.0163 | 0.017 | 0.0308 | 0.0529 | 0.0316 | 0.0293 | 0.0422 | 0.0852 | 0.054 | 0.0571 |  | 0.0374 |
|  | C1 |  | 0.029 | 0.0242 | 0.0212 | 0.0204 | 0.052 | 0.0669 | 0.0278 | 0.0942 | 0.0455 | 0.0711 | 0.0574 |  | 0.0347 |
|  | C2 |  | 0.029 | 0.03 | 0.0217 | 0.0259 | 0.057 | 0.0644 | 0.0281 | 0.0799 | 0.0511 | 0.0656 | 0.0625 |  | 0.0402 |
|  | C3 |  | 0.029 | 0.0242 | 0.0207 | 0.0232 | 0.0629 | 0.0606 | 0.0152 | 0.0814 | 0.0455 | 0.0792 | 0.0563 |  | 0.0406 |
|  | C4 |  | 0.029 | 0.0299 | 0.0225 | 0.03 | 0.0477 | 0.0247 | 0.0125 | 0.0297 | 0.101 | 0.0273 | 0.0319 |  | 0.0284 |
|  | C5 |  | 0.029 | 0.0262 | 0.0207 | 0.0232 | 0.0762 | 0.0758 | 0.0258 | 0.0814 | 0.0455 | 0.0847 | 0.0595 |  | 0.0378 |
|  | D |  |  |  |  | 0.011 |  |  |  |  |  |  |  |  | 0.011 |
|  | E |  | 0.029 | 0.0084 | 0.01 | 0.0277 | 0.0565 | 0.0303 | 0.0091 | 0.0233 | 0.0909 | 0.0437 | 0.0499 |  | 0.0314 |
|  | F |  | 0.0338 | 0.0094 | 0.0069 | 0.0243 | 0.044 | 0.0404 | 0.0101 | 0.0543 | 0.0606 | 0.0219 | 0.0613 |  | 0.03 |
|  | G |  |  |  |  | 0.028 |  |  |  |  |  |  |  |  | 0.028 |
|  | Un |  | 0.029 | 0.0277 | 0.0175 | 0.0229 | 0.0351 | 0.0529 | 0.0122 | 0.0258 | 0.0758 | 0.0437 | 0.0257 |  | 0.0266 |

Supplementary Table-3: Percent nucleotide divergence (PND) and percent amino acid divergence (PAD) between NIV sequences and other genotypes

|  | Genotype | VP1 gene | | Complete Genome | |
| --- | --- | --- | --- | --- | --- |
|  |  | PND | PAI | PND | PAI |
| NIV-C1 vs | A | 16.33% | 96.34% | 19.32% | 95.53% |
|  | B0 | 16.41% | 98.12% | - | - |
|  | B1 | 17.03% | 96.63% | 17.94% | 96.32% |
|  | B2 | 16.34% | 96.65% | 17.87% | 96.10% |
|  | B3 | 16.19% | 97.09% | 18.46% | 96.01% |
|  | B4 | 17.74% | 97.00% | 18.72% | 96.44% |
|  | B5 | 18.30% | 97.18% | 19.35% | 95.82% |
|  | C1 | 6.23% | 99.38% | 7.08% | 99.02% |
|  | C2 | 10.83% | 98.36% | 12.84% | 97.91% |
|  | C3 | 10.55% | 99.28% | 12.57% | 98.44% |
|  | C4 | 12.39% | 97.98% | 17.54% | 96.00% |
|  | C5 | 12.34% | 99.06% | 13.63% | 98.16% |
|  | D | 16.28% | 98.47% | - | - |
|  | E | 17.15% | 96.77% | 17.88% | 96.11% |
|  | F | 18.32% | 96.91% | 19.64% | 95.89% |
|  | G | 15.21% | 95.46% | 15.21% | 95.46% |
|  | Un | 9.61% | 99.32% | 16.36% | 96.94% |
| NIV-D vs | A | 18.09% | 96.36% | 20.04% | 96.57% |
|  | B0 | 17.57% | 98.49% | - | - |
|  | B1 | 17.74% | 96.88% | 17.95% | 96.79% |
|  | B2 | 17.34% | 96.91% | 17.69% | 96.65% |
|  | B3 | 17.59% | 96.52% | 15.16% | 97.67% |
|  | B4 | 17.53% | 96.57% | 17.83% | 96.77% |
|  | B5 | 17.40% | 96.75% | 18.30% | 96.46% |
|  | C1 | 16.55% | 97.81% | 18.63% | 96.70% |
|  | C2 | 17.13% | 97.24% | 19.22% | 96.42% |
|  | C3 | 18.19% | 97.47% | 19.69% | 96.40% |
|  | C4 | 17.18% | 96.87% | 16.88% | 97.38% |
|  | C5 | 18.33% | 97.47% | 19.60% | 96.48% |
|  | D | 5.73% | 98.93% | - | - |
|  | E | 17.09% | 97.02% | 18.26% | 97.21% |
|  | F | 19.04% | 97.13% | 18.98% | 96.98% |
|  | G | 16.23% | 95.75% | - | - |
|  | Un | 16.17% | 97.51% | 17.81% | 97.41% |
| NIV-G vs | A | 19.87% | 97.12% | 20.33% | 96.40% |
|  | B0 | 15.73% | 97.59% | - | - |
|  | B1 | 17.02% | 96.66% | 17.54% | 96.56% |
|  | B2 | 17.13% | 96.67% | 17.68% | 96.44% |
|  | B3 | 16.85% | 96.68% | 15.93% | 97.54% |
|  | B4 | 17.80% | 97.01% | 17.92% | 96.78% |
|  | B5 | 17.09% | 96.92% | 18.31% | 96.26% |
|  | C1 | 16.22% | 97.96% | 18.57% | 96.53% |
|  | C2 | 17.42% | 97.41% | 19.15% | 95.98% |
|  | C3 | 18.34% | 97.68% | 19.73% | 95.94% |
|  | C4 | 17.21% | 97.00% | 17.19% | 97.16% |
|  | C5 | 16.53% | 97.68% | 18.97% | 96.22% |
|  | D | 14.93% | 98.90% | - | - |
|  | E | 16.90% | 97.23% | 17.90% | ret.86% |
|  | F | 17.50% | 97.57% | 18.94% | 97.00% |
|  | G | 10.56% | 97.20% | - | - |
|  | Un | 15.77% | 97.71% | 17.68% | 97.34% |
